# The UAP56 mRNA Export Factor is Required for Dendrite and Synapse Pruning via Actin Regulation in Drosophila

**DOI:** 10.1101/2025.07.23.666284

**Authors:** Samuel Matthew Frommeyer, Ulrike Gigengack, Sandra Rode, Matthew Davies, Sebastian Rumpf

**Affiliations:** Multiscale Imaging Center and Institute for Neurobiology, University of Münster, Röntgenstrasse 16, 48149 Münster, Germany

**Keywords:** dendrite, synapse, pruning, UAP56, actin, cofilin

## Abstract

Neurite and synapse pruning are conserved mechanisms that adapt neuronal circuitry to different developmental stages. Drosophila sensory c4da neurons prune their larval dendrites and their presynaptic terminals during metamorphosis using a gene expression programme that is induced by the steroid hormone ecdysone and involves posttranscriptional regulation pathways. Here we show that loss of the helicase UAP56, an important mediator of nuclear mRNA export, causes strong dendrite and presynapse pruning defects. Loss of UAP56 is linked to actin regulation, as it causes defects in the expression of the actin severing enzyme Mical during dendrite pruning, and actin accumulation at presynapses, where cofilin is required for pruning. Our findings suggest specificity in mRNA export pathways and identify a role for actin disassembly during presynapse pruning.

## Introduction

Neurite and synapse pruning, the regulated degeneration of neuronal connections without loss of the parent neuron, serve to specify neuronal circuits and to remove developmental intermediates during neuronal development (Riccomagno and Kolodkin, 2015, Rumpf et al., 2017). Dysregulation of pruning pathways has been linked to various neurological disorders such as autism spectrum disorders and schizophrenia. The cell biological mechanisms underlying pruning can be studied in Drosophila. For example, the peripheral sensory class IV dendritic arborization (c4da) neurons prune their larval sensory dendrites as well as their presynapses during metamorphosis (Kuo et al., 2005; Williams and Truman, 2005, Furusawa et al., 2023). Dendrite pruning is induced by an ecdysone pulse at the onset of metamorphosis that induces the expression of pruning factors such as the transcription factor Sox14 and the actin severing enzyme Mical (Kirilly et al., 2009) and also leads to changes in neuronal metabolism (Marzano et al., 2021). A key target during large scale pruning is the microtubule cytoskeleton, which is locally disassembled in dendrites (Herzmann et al., 2017; Herzmann et al., 2018; Rumpf et al., 2019, Davies et al., 2025 preprint), eventually leading to mechanical tearing of the whole dendrite close to the cell body (Krämer et al., 2023). While c4da neuron axons stay intact during metamorphosis (Kuo et al., 2005), their presynaptic terminals are also pruned in an ecdysone-dependent manner, but few downstream pathways are known (Furusawa et al., 2023).

As c4da neuron pruning is induced by changes in gene expression, several post-transcriptional control mechanisms have been implicated in pruning as well, including splicing (Rumpf et al., 2014; Rode et al., 2017) and translation control (Rode et al., 2018). Specific requirements in the translation machinery during neuronal remodeling could be linked to differential activity of the Target of Rapamycin (TOR) pathway during development (Wong et al., 2013; Sanal et al., 2023).

Nuclear mRNA export occurs after - or concomiantly with - splicing. Here, a set of dedicated export factors including the DEXD box ATPase UAP56 (also known as HEL25E in Drosophila), the THO complex and the protein Aly/Ref1 are co-transcriptionally loaded onto nascent mRNP complexes. This UAP56/THO/Aly complex is then disassembled at the nuclear pore by the TREX complex, making the mRNA export-competent (Xie and Ren, 2019). While these factors are thought to be involved broadly in the export of most or all mRNAs, there is also some evidence for specificity in mRNA export pathways. In neurons, for example, the THO complex was shown to be particularly important for export of mRNAs related to synapses and dopamine (Maeder et al., 2018). In support of specific nervous system-related function(s) for UAP56 also in humans, congenital mutations in the UAP56 homologue DDX39B were recently shown to cause neurodevelopmental defects (Booth et al., 2025).

Here, we show that UAP56 is required for c4da neuron dendrite pruning. While several pruning pathways are not strongly affected by loss of UAP56, cytoplasmic levels of *MICAL* mRNA are reduced, and Mical protein expression is delayed, suggesting that *MICAL* mRNA may be strongly dependent on UAP56. Furthermore, we find that UAP56 loss causes presynapse pruning defects that also involve defects in actin remodeling. In contrast to dendrites, where Mical is important, actin disassembly at the presynapse seems to depend more strongly on the actin disassembly factor cofilin. Thus, our results describe a specific neurodevelopmental function for an mRNA export factor involved in human neurological disease.

## Results

### The mRNA export factor UAP56 is required for c4da neuron dendrite pruning

C4da neurons prune their larval dendrites within the first 18 hours after puparium formation (h APF) (Fig. 1 A, B, B’). To gain further insight in posttranscriptional regulation of dendrite pruning, we conducted an RNAi screen for mRNA-related factors. We used the c4da-specific driver *ppk-GAL4* to express dsRNA lines against 92 genes that caused splicing defects in S2 cells (Park et al., 2004, Rode et al., 2018) and identified a line directed against UAP56/HEL25E. C4da neurons expressing UAP56 dsRNA exhibited relatively normal dendrites at the third instar stage (Figure 1 C) but displayed strong dendrite pruning defects at 18 h APF (Figure 1 C’, F, G). We confirmed these results using a second dsRNA line directed against UAP56 (Figure 1 D - D’, F, G) which also caused strong dendrite pruning defects. To verify these results by an independent method, we performed Mosaic Analysis with a Repressible Marker (MARCM) with *uap56^k11511^*, a lethal P element insertion in the 5’ untranslated region of the UAP56 locus. C4da neurons homozygous mutant for *uap56* also displayed strong dendrite pruning defects at 18 h APF (Figure 1 E - G).

**Figure 1.**
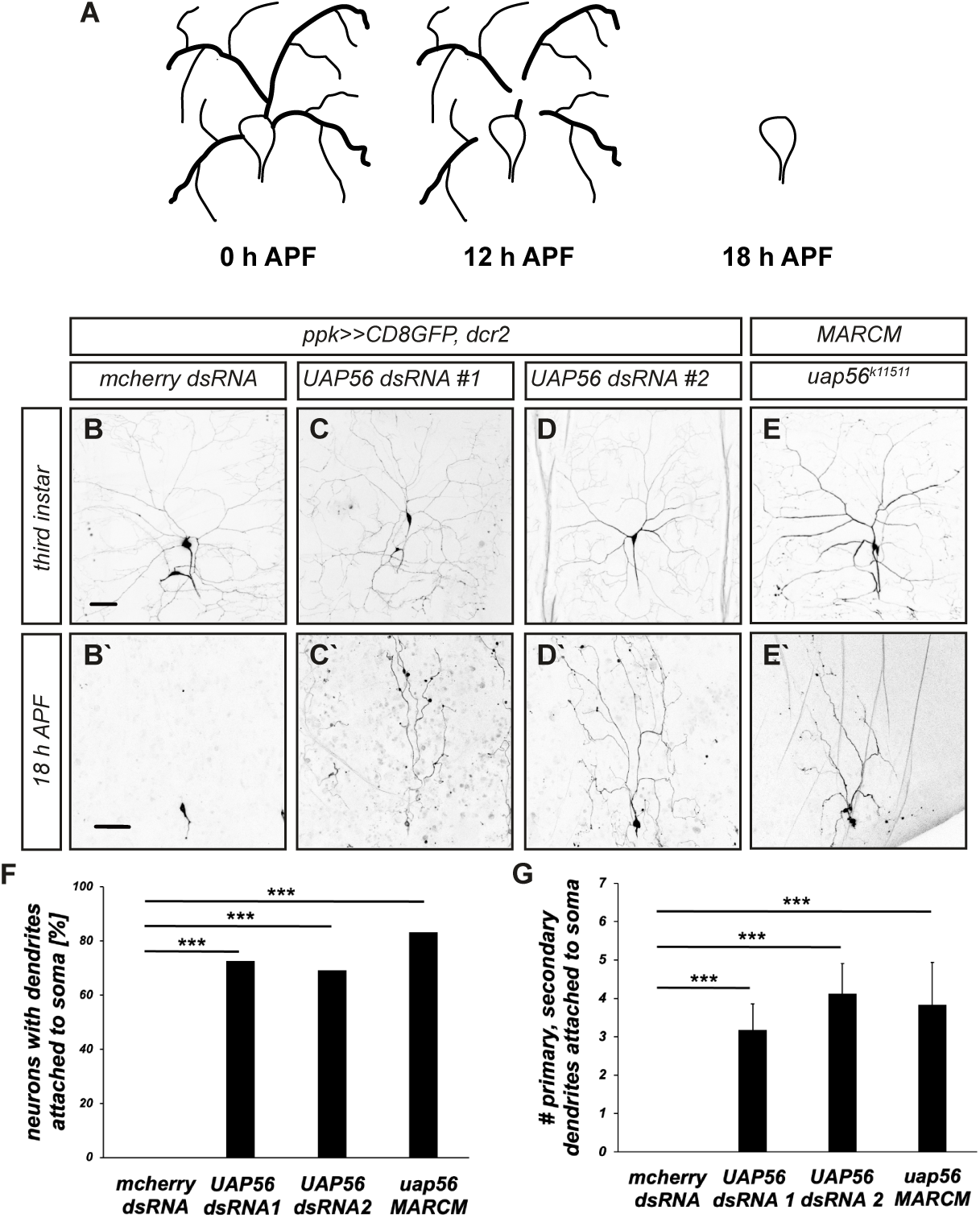
UAP56 is required for c4da neuron dendrite pruning. **A** Schematic timecourse of c4da neuron dendrite pruning. **B** – **E’** C4da neurons were labeled by CD8::GFP expression under *ppk-GAL4* or tdtomato expression in MARCM clones and visualized at the third instar larval stage (**B - E**) or at 18 h APF (**B’ - E’**). **B, B’** Control c4da neurons expressing mcherry dsRNA. **C, C’** C4da neurons expressing UAP56 dsRNA#1. **D, D’** C4da neurons expressing UAP56 dsRNA#2. **E, E’** Homozygous *uap56^k11511^* c4da neuron MARCM clones. **F** Penetrance of pruning defects in **B’**- **E’**. N=24-25, *** P<0.0005, two-tailed Fisher’s exact test. **G** Severity of pruning defects as assessed by the number of attached primary and secondary dendrites in **B’**- **E’**. N=24- 25, *** P<0.0005, Wilcoxon’s test. Scale bars in **B**, **B’** are 50 µm.

### Role of the UAP56 ATPase

UAP56 is an ATP-dependent helicase involved in mRNA export from the nucleus. Its ATPase activity is thought to be required for the disassembly of a mRNA export complex containing the THO complex and Aly/Ref1 at the nuclear pore (Xie and Ren, 2019). To study the function of UAP56 in c4da neurons, we generated C-terminally GFP-tagged UAP56 transgenes for use with the UAS system. UAP56::GFP mainly localized to the nucleus and nuclear periphery in c4da neurons (Figure 2 A - A’’). To assess a potential function of the ATPase, we next disrupted the conserved DECD motif in the UAP56 ATPase domain by mutating the glutamate in the motif (at position 194) to glutamine. In contrast to wild type UAP56::GFP, the UAP56::GFP^E194Q^ variant was evenly distributed in the c4da neuron nucleus and cytoplasm (Figure 2 B - B’’). To test whether the ATPase affects UAP56 protein interactions, we performed co-immunoprecipitations with the known UAP56 binding partner Ref1 in S2 cells. While myc-tagged Ref1 could not be detected in precipitates of wild type UAP56::GFP, it was clearly detectable in UAP56::GFP^E194Q^ precipitates (Figure 2 C), suggesting that an active UAP56 ATPase may weaken or disassemble the UAP56/Ref1 complex. To assess whether the ATPase is necessary for dendrite pruning, we next expressed wild type UAP56::GFP or UAP56::GFP E194Q in *uap56^k11511^*MARCM clones and tested whether they can rescue the pruning defects. Surprisingly, both wild type UAP56 and the E194Q mutant efficiently rescued the pruning defects of the mutant c4da neurons at 18 h APF (Figure 2D - F’, G, H). Thus, our data suggest that UAP56 ATPase activity is required for its localization and disassembly of a UAP56/Ref1 complex, but not stringently for its function during c4da neuron dendrite pruning.

**Figure 2.**
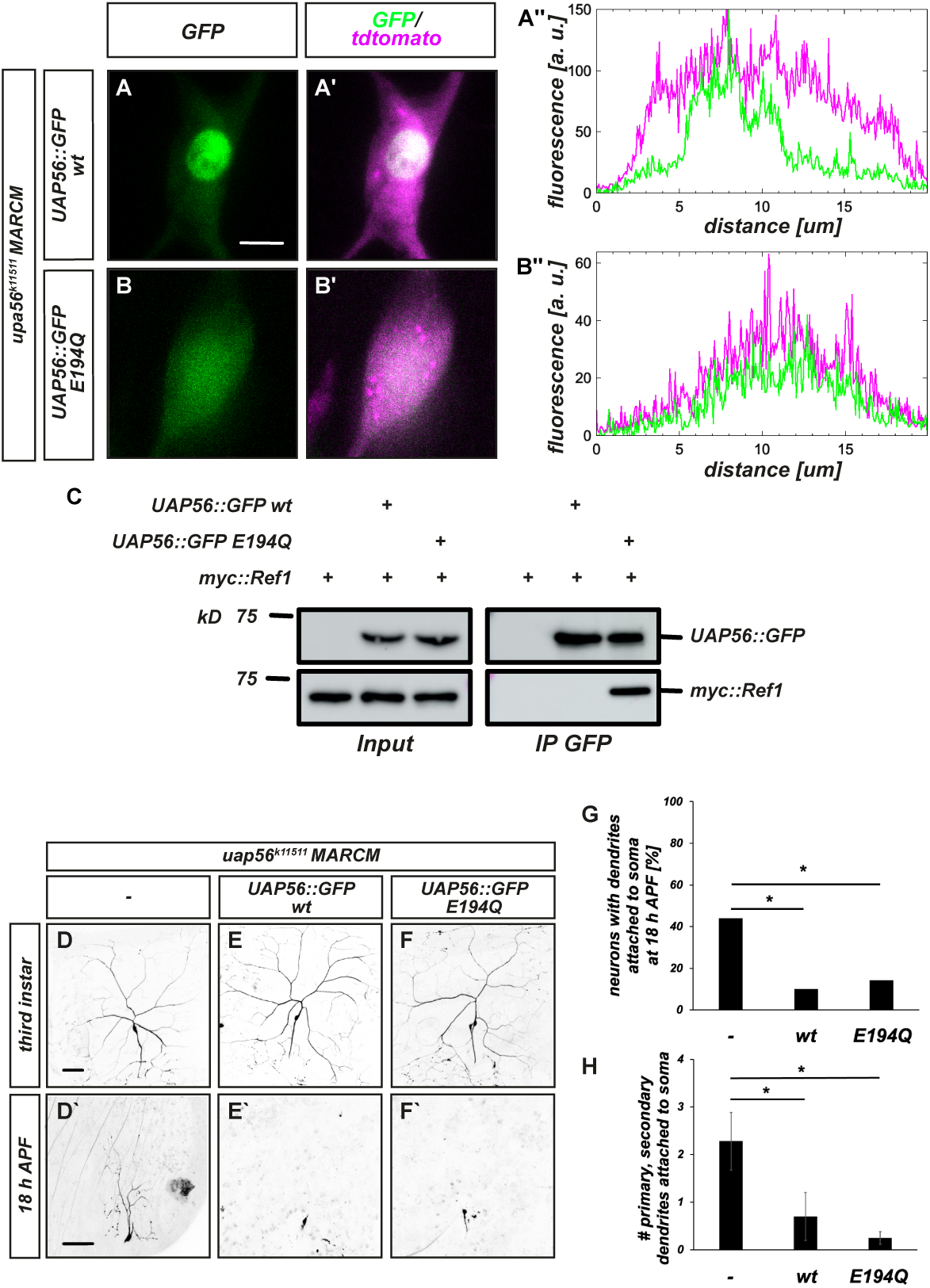
The ATPase affects UAP56 localization, but not c4da neuron dendrite pruning. **A** - **B’’** Localization of GFP-tagged UAP56 wt (**A** - **A’’**) or UAP56 E194Q (**B** - **B’’**). The indicated UAS transgenes were expressed in *uap56* MARCM clones and visualized by confocal microscopy. Panels **A**, **B** show GFP signal, panels **A’**, **B’** show merge with cytosolic tdtomato, and panels **A’’**, **B’’** show intensity profiles across cell body and nucleus. **C** Co-immunoprecipitation between UAP56 and Ref1. GFP-tagged UAP56 transgenes (wt or E194Q) were coexpressed with myc-Ref1 in S2 cells, and precipitated using anti-GFP antibodies. Input (left) and bound fractions (right) are shown, the position of a molecular weight marker is indicated to the left. **D** – **H** Rescue experiments with UAP56 wt or E194Q. *uap56^k11511^* c4da neuron MARCM clones are shown at the third instar (**D** - **F**) or at 18 h APF (**D’** - **F’**). **D**, **D’** *uap56^k11511^* c4da neuron MARCM clones. **E**, **E’** *uap56^k11511^* c4da neuron MARCM clones expressing UAP56::GFP wt. **F**, **F’** *uap56^k11511^*c4da neuron MARCM clones expressing UAP56::GFP E194Q. **G** Penetrance of c4da neuron dendrite pruning defects in **D’** - **F’**. N= 18 - 20, * P<0.05, Fisher’s exact test. **H** Severity of c4da neuron dendrite pruning defects as assessed by the number of attached primary and secondary dendrites in **D’** - **F’**. N= 18 - 20, * P<0.05, Wilcoxon’s test. Scale bars in **A**, **D** and **D’** are 10 and 50 µm, respectively.

### Loss of UAP56 does not affect microtubule regulation or the ubiquitin system

To begin to address how UAP56 deficiency affects c4da neuron dendrite pruning, we next assessed the effects of UAP56 knockdown on various known pruning pathways. C4da neuron dendrite pruning requires organized microtubule disassembly that depends on proper plus end-in orientation of dendritic microtubules (Rumpf et al., 2019). To test for normal organization of the dendritic microtubule cytoskeleton, we used live imaging of EB1::GFP. Neither the speed nor the directionality of EB1::GFP comets were altered in neurons expressing UAP56 dsRNA when compared to neurons expressing a control dsRNA against the odorant receptor Orco, indicating largely normal microtubule organization in neurons lacking UAP56 (Figure 3 A - C).

**Figure 3.**
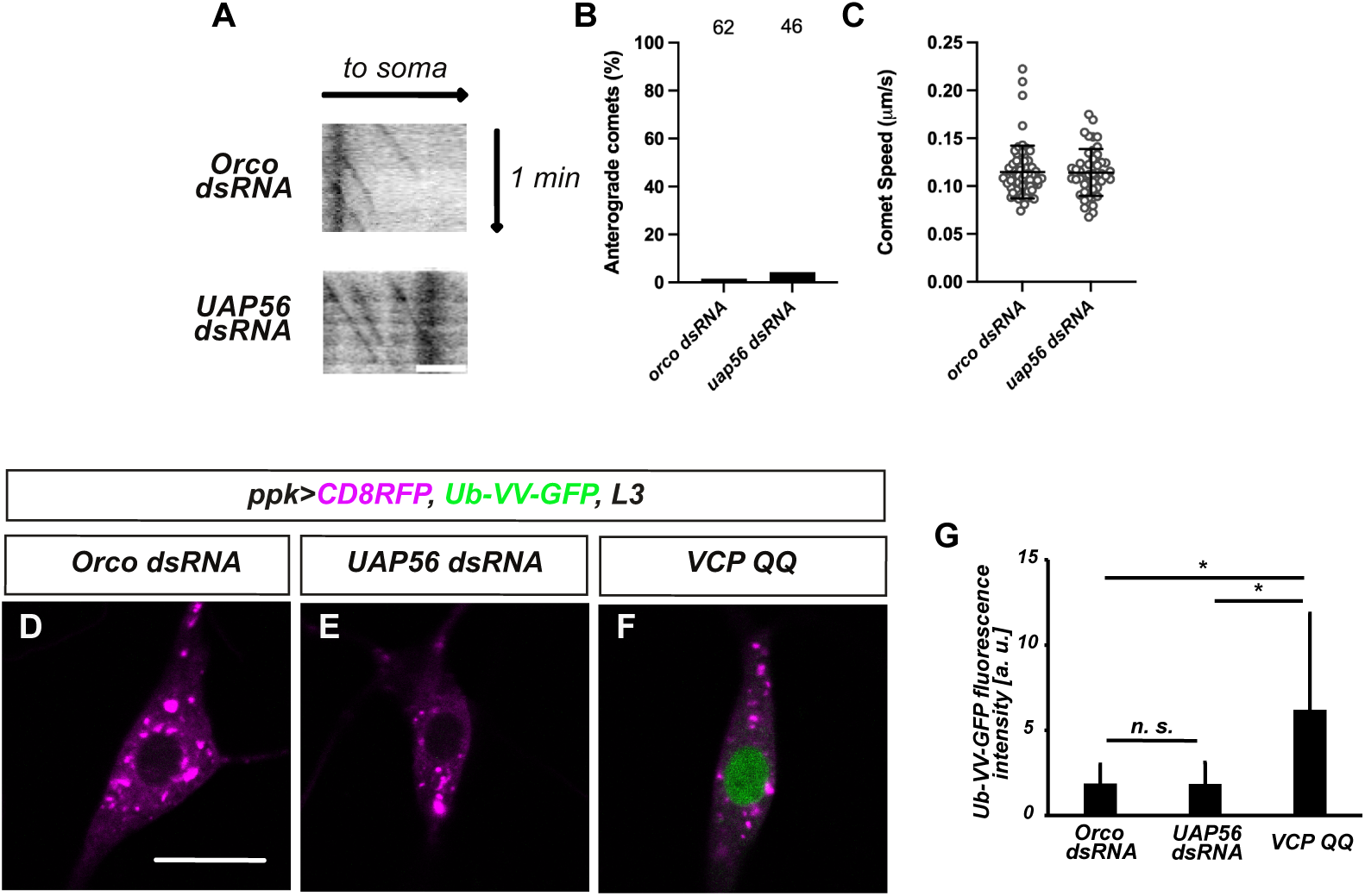
UAP56 loss does not affect microtubule organization or ubiquitin- mediated protein degradation. **A** – **C** Microtubule orientation and growth speed were determined using EB1::GFP. **A** Dendritic EB1::GFP kymographs from control c4da neurons expressing Orco dsRNA (upper panel) or UAP56 dsRNA (lower panel). Upper arrow indicates direction of soma (retrograde). **B** Percentage of anterograde comets in **A**, N is indicated in the graph. Fisher’s exact test P value was not significant. **C** Comet speed in **A**. N=62 and 46, respectively, the average is indicated by a line. Whiskers represent standard deviation, the Wilcoxon’s test P value was not significant. **D** – **G** Use of a Ub-VV-GFP reporter to assess gene loss effects on ubiquitin-proteasome system function in c4da neurons. Panels **D** – **F** show reporter GFP fluorescence together with CD8::RFP to label the neuron. **D** Control c4da neuron expressing Orco dsRNA. **E** C4da neuron expressing UAP56 dsRNA. **F** C4da neuron expressing dominant-negative VCP QQ. **G** Quantification of Ub-VV-GFP fluorescence intensity in panels **D** - **F**. N=9-11, values are mean +/- s. d., * p<0.05, Student’s t-test. Scale bars in **A** and **D** are 5 μm and 10 μm, respectively.

C4da neuron dendrite pruning also requires the ubiquitin-proteasome system (UPS) (Kuo et al., 2005; Rumpf et al., 2011; Wong et al., 2013). To assess global UPS function, we used a Ub-VV-GFP reporter gene, a ubiquitin fusion degradation (UFD) substrate. Here, ubiquitin is fused to the N-terminus of GFP, and the C-terminal glycine residues of ubiquitin are changed to valine, rendering the fusion uncleavable by deubiquitinating enzymes and thus destabilizing it. Ub-VV-GFP was undetectable in c4da neurons expressing both control and UAP56 dsRNAs (Figure 3 D, E, G). As a proof of reporter functionality, we expressed VCP QQ, a dominant-negative version of the UPS chaperone VCP (Rumpf et al., 2011) which caused stabilization of the Ub-VV-GFP reporter (Figure 3 F, G). Thus, several important cellular pruning pathways are not affected by loss of UAP56.

### Loss of UAP56 affects Mical expression

The ecdysone/Sox14/MICAL pathway is an important transcriptional pathway during dendrite pruning (Kirilly et al., 2009). Sox14 could be detected in c4da neurons expressing UAP56 dsRNA at both 0 and 2 h APF (Figure S1). The Sox14 target gene in the dendrite pruning pathway, *MICAL*, encodes a large actin severing protein of 4723 amino acids (from a 15 kb mRNA). We therefore reasoned that Mical expression could be more challenging to c4da neurons. To test whether loss of UAP56 affects *MICAL*, we next visualized Mical mRNA in c4da neurons using the MS2/MCP system (Bertrand et al., 1998). To this end, we introduced six copies of the MS2 sequence into the Mical 3’ untranslated region (3’UTR) using a CRISPR-based strategy (Zhang et al., 2014) (Fig. 4 A). We then expressed an RFP-tagged version of the MS2 binding protein MCP in c4da neurons. MCP contains a nuclear localization signal, and is mainly localized in the nucleus, except when it is exported as part of an MS2-tagged mRNA particle. Larval c4da neurons, which do not express Mical, contained approximately ten cytoplasmic MCP::RFP dots irrespective of background (control or *Mical*-MS2_6_), likely due to strong MCP::RFP expression under *ppk-GAL4* at this stage (Fig. S2). At 2 h APF, however, control c4da neurons displayed only 3 - 4 dim puncta in the cytosol (Figure 4 B, E). In contrast, on average eight bright MCP::RFP puncta could be seen in homozygous *Mical*-MS2_6_ animals at this stage (Figure 4 C, E). To ascertain that these dots represented tagged *Mical* mRNA, we expressed EcR DN, a dominant-negative version of the ecdysone receptor. This reduced the number of MCP::RFP puncta to background levels (Figure 4 D, E). We next expressed UAP56 dsRNA in c4da neurons and found that this strongly reduced the number of cytoplasmic MCP::RFP puncta (Figure 4 F - H), indicating that Mical mRNA depends on UAP56 for its nuclear export. To verify this notion, we next assessed the effect of UAP56 knockdown on Mical protein expression by immunofluorescence. Mical expression in control neurons is detectable as early as the white pupal stage (0 h APF) and persists beyond 2 h APF (Figure 5 A, B, E). Consistent with the reduced amount of Mical mRNA upon UAP56 knockdown, Mical protein could not be detected at 0 h APF (Figure 5 C, E). However, we detected almost normal levels of Mical protein at 2 h APF (Figure 5 D, E), indicating that UAP56 knockdown leads to a delay in Mical protein accumulation. Unfortunately, co-expression of UAP56 dsRNA constructs with other UAS constructs led to weaker effects of the dsRNA, likely via titration, so we could not reliably assess whether exogenous Mical expression can rescue the UAP56 loss of function phenotype. However, loss of Mical expression is at least partially causal to the pruning defects of a number of other mutants (Kirilly et al., 2009; Rode et al., 2018; Wolterhoff et al., 2020), indicating that UAP56 deficiency affects c4da neuron dendrite pruning at least in part by delaying Mical expression.

**Figure 4.**
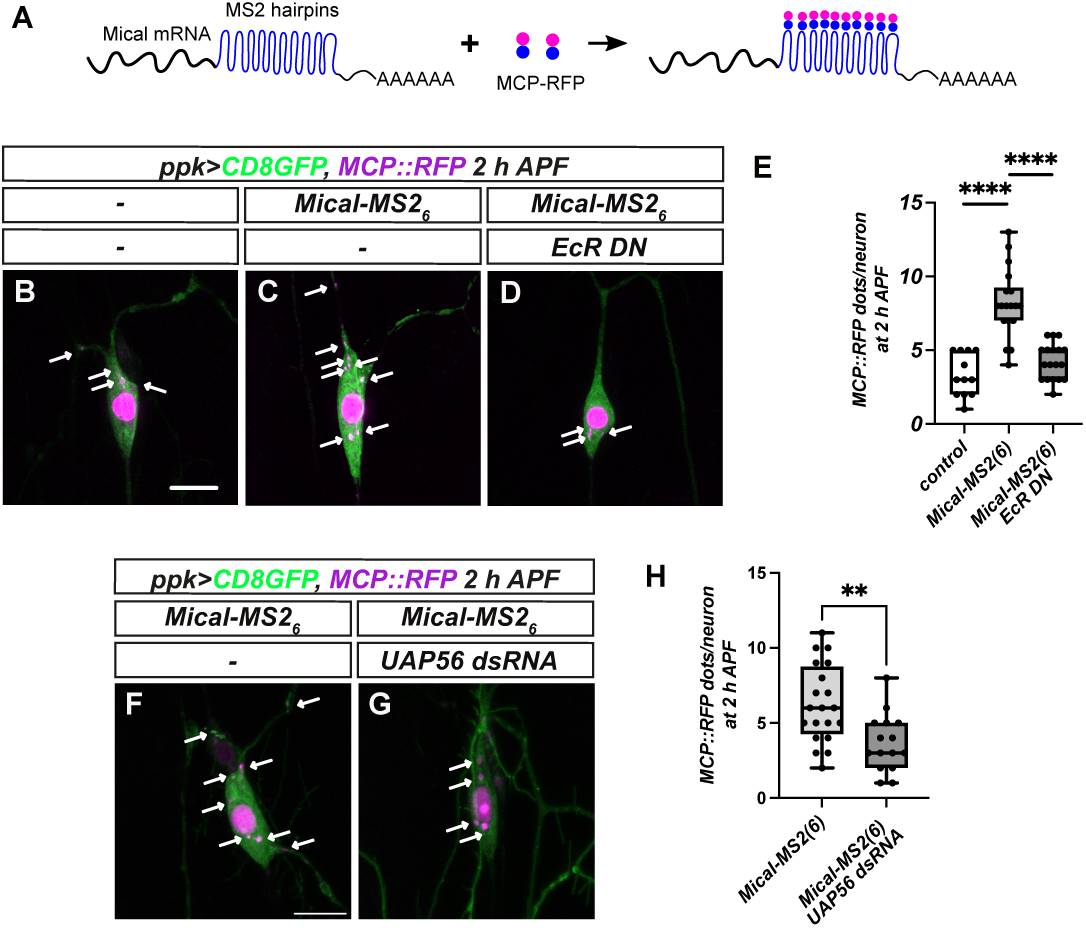
Establishment of a *Mical* mRNA reporter and effect of UAP56. **A** *Mical* mRNA was labeled by knock-in of the MS2 sequence in the 3’UTR, enabling visualization by expression of MCP::RFP. **B** - **E** Characterization of the *MICAL-MS2_6_* reporter. MCP::RFP was coexpressed with CD8::GFP in c4da neurons and visualized at 2 h APF. Cytoplasmic MCP::RFP puncta are labeled by arrows. **B** Control c4da neuron in a wild type background. **C** Control c4da neuron in the *MICAL-MS2_6_* background. **D** C4da neuron expressing dominant-negative EcR-DN in the *MICAL- MS2_6_* background. **E** Number of cytoplasmic MCP::RFP puncta in **B** - **D**. Boxes represent first quartiles and the median is indicated. Whiskers show standard deviation, N=12 - 18, *** P<0.0005, Wilcoxon’s test. **F** - **H** Effect of UAP56 knockdown on *MICAL-MS2_6_* at 2 h APF. **F** Control c4da neuron in the *MICAL-MS2_6_* background. **G** C4da neuron expressing UAP56 dsRNA in the *MICAL-MS2_6_* background. **H** Quantification of cytoplasmic MCP::RFP puncta in **F**, **G**. N=20 and 15, respectively. ** P<0.005, Wilcoxon’s test. Scale bars in **B** and **F** are 10 μm.

**Figure 5.**
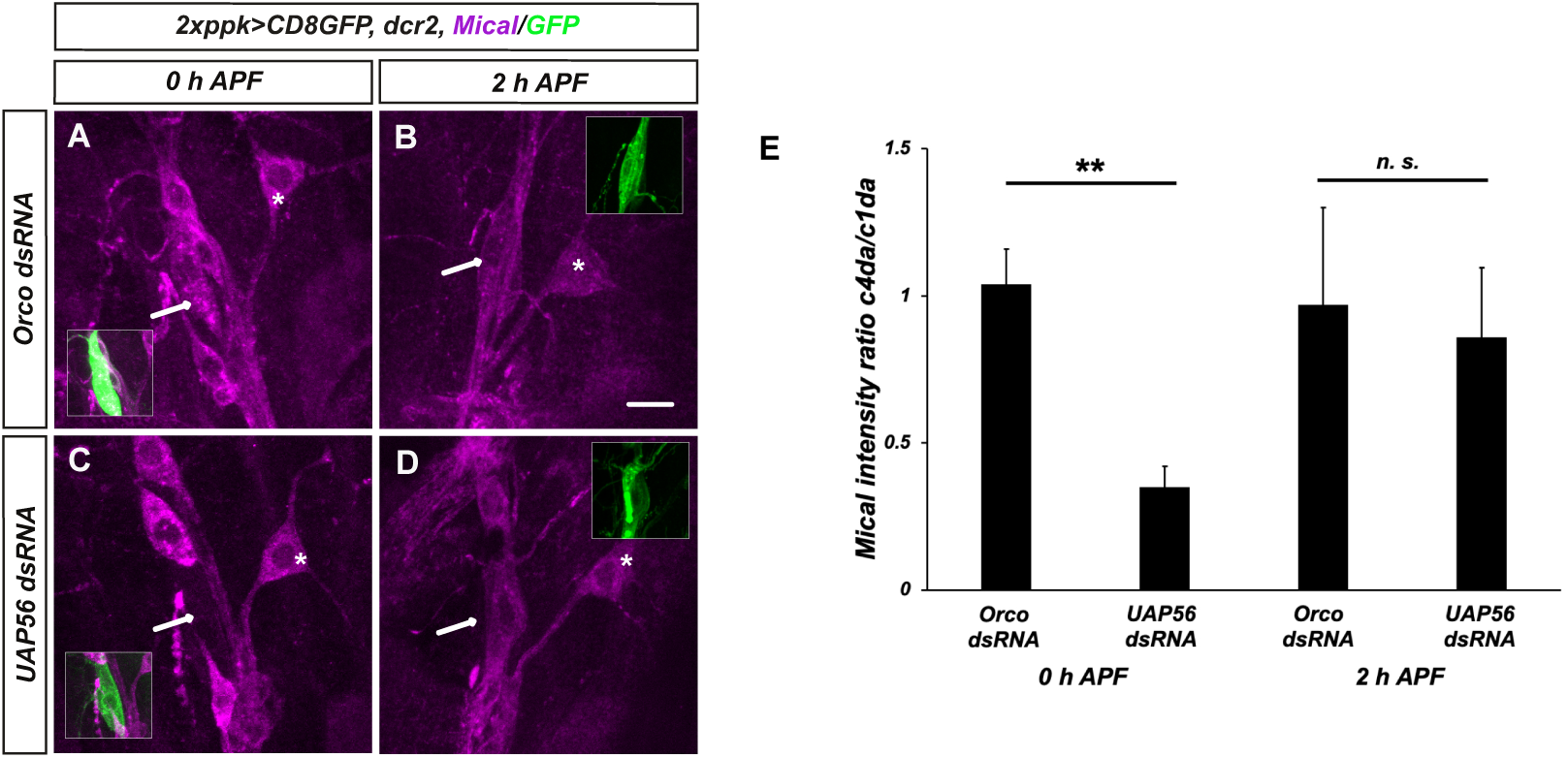
UAP56 loss causes a delay in Mical protein expression. **A** - **D** Mical was visualized by immunofluorescence at the 0 h APF (white pupal stage) or at 2 h APF. The indicated dsRNAs and GFP (insets) were driven in c4da neurons by *ppk-GAL4*. **A** Control c4da neuron expressing a dsRNA construct against Orco at 0 h APF. **B** Control c4da neuron at 2 h APF. **C** C4da neuron expressing a dsRNA construct against UAP56 at 0 h APF. **D** C4da neuron expressing a dsRNA construct against UAP56 at 2 h APF. **E** Quantification of Mical expression levels in **A** - **D**. Mical expression levels in c4da neurons (arrows in **A** - **D**) were normalized to expression levels in unaffected neighboring c1da neurons (asterisks). N=6 (0 h APF) or N=8-9 (2 h APF). Values are mean +/- s. d., n. s., not significant, ** P<0.005, Student’s t-test. The scale bar in **B** is 10 μm.

### UAP56 is required for c4da neuron presynapse pruning

C4da neuron axons project to the sensory domain of the ventral nerve cord (VNC), where they form segmental commissures across the midline that contain presynaptic active zones characterized by the ELKS/CAST protein Bruchpilot (Brp) (Fig 6 A). While c4da neuron axons persist during metamorphosis, these commissures and presynapses are pruned in an ecdysone- and ubiquitin-dependent manner with a similar timecourse as c4da neuron dendrites (Furusawa et al., 2023) (Fig. 6 E). We next assessed whether UAP56 knockdown also caused presynapse pruning defects. To specifically visualize c4da neuron presynapses, we expressed a fluorescently tagged Brp fragment, Brp^short^::Strawberry, under *ppk-GAL4*. In control animals at the larval stage, approximately 400 presynaptic Brp^short^::Strawberry puncta could be seen in each segment (Fig. 6 B, D), and UAP56 knockdown did not affect their number or distribution (Fig. 6 C, D). At 24 h APF, c4da neuron axonal commissures in segments 3 - 5 in control neurons had disappeared, and the number of presynaptic Brp^short^::Strawberry puncta was strongly reduced to approximately 20 per segment (Fig. 6 F, H). Furthermore, more diffuse red fluorescent signal could sometimes be observed that did not overlap with the persisting axon structures, possibly indicating uptake of presynaptic material into surrounding tissues (Fig. 6 F). In UAP56 knockdown c4da neurons, axonal commissures were frequently still intact, and approximately 90 Brp^short^::Strawberry puncta could be detected in each segment (P<0.001, student’s t-test) (Fig. 6 G, H), indicative of synapse pruning defects. Thus, loss of UAP56 causes widespread defects in neuronal remodeling.

**Figure 6.**
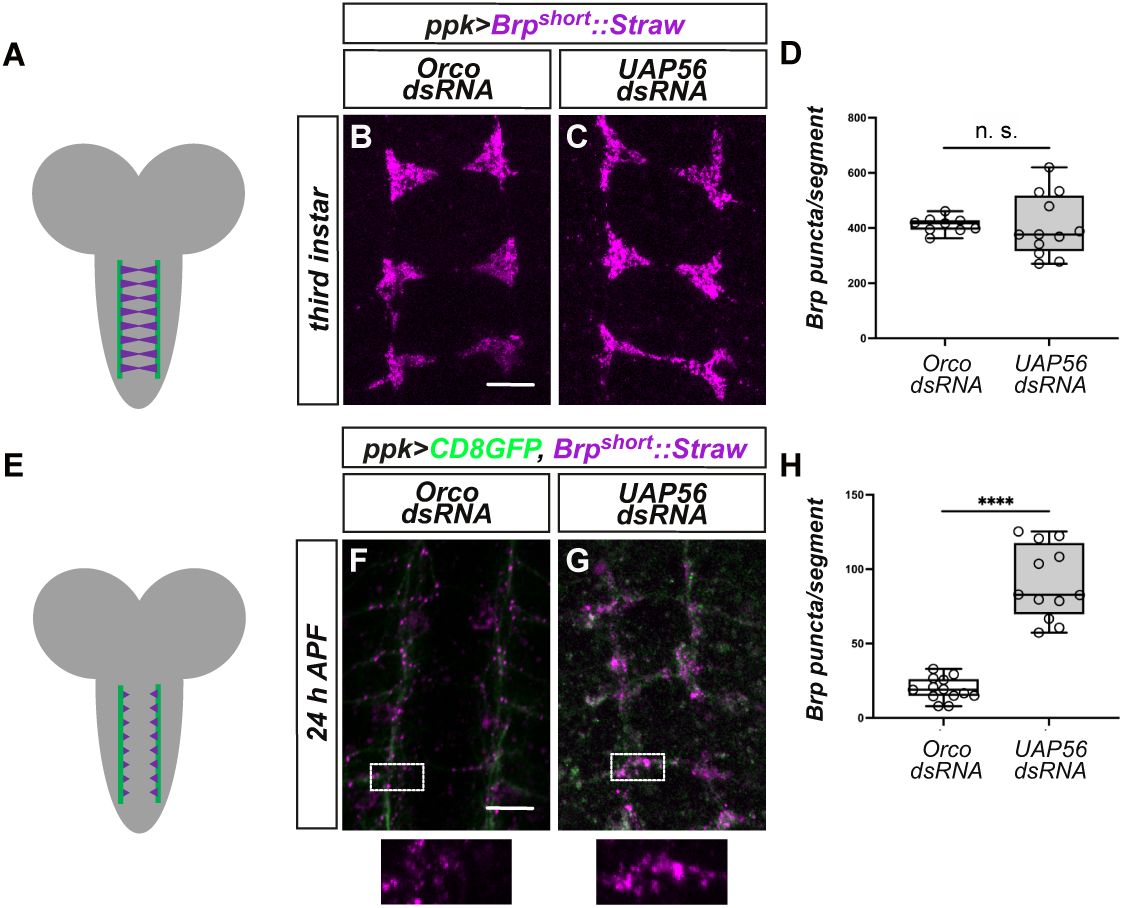
Loss of UAP56 causes c4da neuron presynapse pruning defects. **A** Schematic of the ladder-like segmental arrangements of c4da neuron axons (green) and presynapses (magenta) in the larval VNC. **B** - **D** Effect of UAP56 loss on c4da neuron active zone number in third instar larvae. C4da neuron active zones in the VNC were labeled by expression of *Brp^short^::Strawberry* via *ppk-GAL4*. Shown are segments 3-5. **B** Control c4da neurons expressing Orco dsRNA. **C** C4da neurons expressing UAP56 dsRNA. **D** Number of *Brp^short^::Strawberry* puncta in larvae. The average number of synaptic Brp puncta per segment was determined for each animal using SynQuant. Box indicates inner quartiles, the median is indicated by a line. Whiskers represent standard deviation. N = 9, 13 animals, n. s., not significant, Student’s t-test. **E** Schematic of c4da neuron axon morphology at 24 h APF. **F** - **H** C4da neuron active zones at 24 h APF. Lower panels show magnified views of the *Brp^short^::Strawberry* signal in the boxed areas above. **F** Control animals expressing Orco dsRNA in c4da neurons. **G** Animals expressing UAP56 dsRNA. **H** Number of *Brp^short^::Strawberry* puncta at 24 h APF. N = 14, 12, *** P<0.0005, Student’s t-test. Scale bars are 10 μm in **A** and **D**.

### Presynapse pruning requires actin remodeling

As loss of UAP56 affects expression of the actin-severing enzyme Mical during c4da neuron dendrite pruning, we next asked whether defects in actin remodeling could also underlie the effects of UAP56 loss at presynapses. To address this, we expressed the actin marker lifeact::GFP in c4da neurons and visualized it in the VNC. At 24 h APF, when axonal commissures have been pruned, lifeact::GFP still evenly labeled the ipsilateral axon endings in control neurons (Fig. 7 A). UAP56 knockdown led to persistence of c4da neuron axonal commissures, which were clearly labeled by lifeact::GFP, and the marker was also visible in blob-like structures at synaptic sites (Fig. 7 B), indicating that defective actin disassembly may indeed contribute to c4da neuron presynapse pruning defects upon UAP56 knockdown.

**Figure 7.**
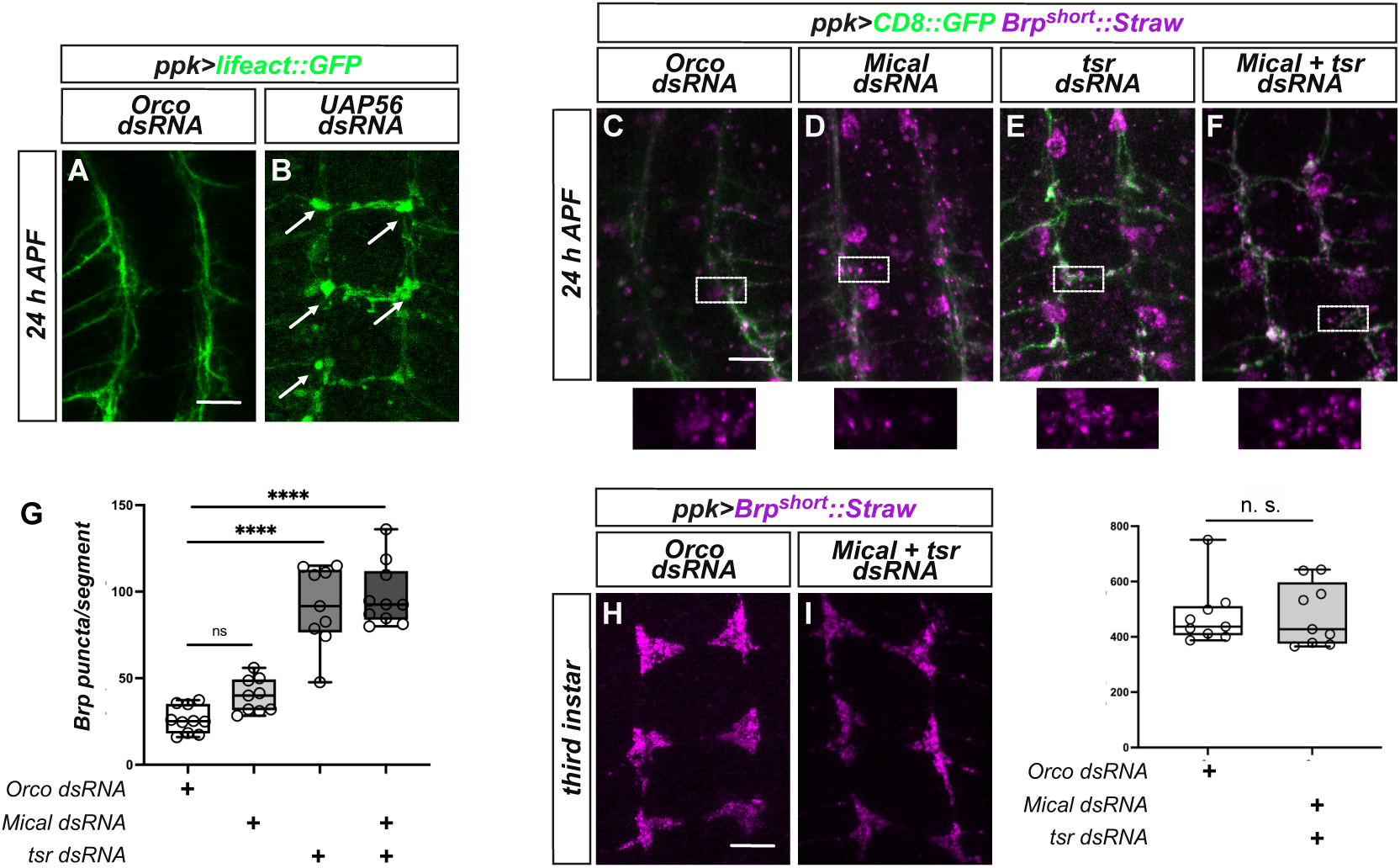
Actin disassembly factors are required for c4da neuron presynapse pruning. **A**, **B** Loss of UAP56 causes actin accumulation at presynaptic sites during the pupal stage. F-actin was labeled with *UAS-lifeact::GFP* under *ppk-GAL4*, and c4da neuron axonal endings in the VNC in segments 3 - 5 were visualized at 24 h APF. **A** Control c4da neurons expressing Orco dsRNA. **B** C4da neurons expressing UAP56 dsRNA. Actin accumulations are indicated by arrows. **C** - **G** Actin disassembly factors are required for c4da neuron presynapse pruning. C4da neuron axons and presynapses in the VNC were labeled by expression of CD8::GFP and the active zone marker *Brp^short^::Strawberry* under the control of *ppk-GAL4*, and VNCs were visualized at 24 h APF. Lower panels show magnified views of the *Brp^short^::Strawberry* signal in the boxed areas above. **C** Control neurons expressing Orco dsRNA. **D** Neurons expressing Mical dsRNA. **E** Neurons expressing tsr dsRNA. **F** Neurons expressing both Mical and tsr dsRNA. **G** Quantification of the number of Brp puncta in **C** - **F**. Analysis was done as in Figure 6. N=9 - 10, n. s., not significant, *** P<0.0005, two-way ANOVA. **H** - **J** Actin disassembly does not affect c4da neuron active zone number at the larval stage. Active zones labeled by *Brp^short^::Strawberry* were visualized at the third instar stage. **H** C4da neuron active zones in control larvae expressing Orco dsRNA. **I** C4da neuron active zones at 24 h APF in larvae expressing both Mical and tsr dsRNA. **J** Quantification of the number of Brp puncta in **H**, **I**. N=9 each, n. s., not significant, student’s t-test. Scale bars are 10 μm in **A**, **C** and **H**.

To further verify the role of actin disassembly during synapse pruning, we asked whether actin disassembly by Mical is also required for c4da neuron presynapse pruning. At 24 h APF, Mical knockdown alone increased Brp puncta only mildly and non-significantly compared to a control (Fig. 7 C, D, G). Actin can not only be disassembled through oxidation by Mical, but also through depolymerization from minus ends or through severing, usually performed by the actin depolymerizing factor cofilin. We therefore next tested cofilin, encoded by *twinstar* (tsr) in Drosophila, and found that tsr knockdown caused strong synapse pruning defects (Fig. 7 E, G). Cofilin and Mical can act synergistically, especially on bundled actin fibers (Rajan et al., 2023), but this is unlikely to be the case during presynapse pruning as coexpression of dsRNA constructs against Mical and *tsr* did not significantly increase the number of remaining Brp^short^::Strawberry puncta compared to *tsr* knockdown alone (Fig. 7 F, G). Importantly, the observed effects were specific for synapse pruning during metamorphosis, as combined knockdown of Mical and tsr did not significantly alter the number of presynaptic Brp^short^::Strawberry puncta at the larval stage (Fig. 7 H - J). Taken together, our data indicate that loss of UAP56 affects actin remodeling during c4da neuron presynapse pruning, a process that normally requires cofilin.

## Discussion

Here, we describe a role for the mRNA export factor UAP56 during c4da neuron neuronal remodeling. While a general inhibition of mRNA export might be expected to cause broad defects in cellular processes, our data in develoing c4da neurons indicate that loss of UAP56 mainly affects pruning mechanisms related to actin remodeling. Firstly, our data suggest that UAP56 affects expression of the actin severing enzyme Mical. Mical mRNA is downregulated in the cytosol upon loss of UAP56, suggesting that it is a direct UAP56 target. In addition to dendrite pruning, UAP56 is also required for presynapse pruning. Here, the actin marker lifeact::GFP accumulates at presynaptic sites during pruning upon UAP56 knockdown, suggesting that UAP56 may also affect actin in presynapses. Indeed, we found that presynapse pruning also depends on actin disassembly, this time not by Mical, but by cofilin. Whether cofilin is also a UAP56 target remains to be seen, alternatively, several cofilin upstream regulators have been described (Rust, 2015) which might also be involved in presynapse pruning and subject to UAP56 regulation. Thus, our data suggest that UAP56 is particularly important for the export of a set of functionally related mRNAs in c4da neurons, further supporting the idea that mRNA export factors might have specific targets.

Cofilin has previously been implicated in both pre- and postsynaptic processes such as dendritic spine maturation, long-term depression, and negative regulation of neurotransmitter release (Rust, 2015), but not in pruning. Our data also reveal a surprising specialization in actin remodeling factors: Whereas Mical is the predominant actin severing enzyme during c4da neuron dendrite pruning, cofilin is more important during synapse pruning. It is interesting to speculate that distinct actin structures must be disassembled in these compartments.

In our experiments, we also addressed the role of the UAP56 ATPase. UAP56 E194Q, a mutant predicted to inhibit or slow down UAP56 ATPase activity, affected UAP56 localization and protein interactions, but it efficiently rescued the pruning defects of a *uap56* mutant allele. One explanation for this could be that UAP56 does not need to be removed from mRNP particles during nuclear export for them to be translated. Since our mutant allele *uap56^k11511^* is a P element insertion in the 5’UTR and therefore likely a (strong) hypomorph, the rescue could also point to a cooperative mechanism of action, where residual wild type UAP56 together with UAP56 E194Q may be able to support the cellular needs. Taken together, our data provide new insights into the function of posttranscriptional gene regulation during neuronal development.

## Materials and Methods

### Fly Strains

For expression in c4da neurons, we used *ppk-GAL4* insertions on the second and third chromosomes (Grueber et al., 2007) driving UAS-CD8::GFP. MARCM clones were induced with *SOP-FLP* (Matsubara et al., 2011) and labeled by UAS-tdtomato (Han et al., 2014) expression under *nsyb-GAL4^R57C10^* (Pfeiffer et al., 2008). Other UAS lines were UAS-EcR^DN^ (BL 6872), UAS-MCP::RFP (BL 27418), UAS-VCP QQ (Rumpf et al., 2011), UAS-lifeact::GFP (BL 35544), UAS- Brp^short^::Strawberry (gift from S. Sigrist). dsRNA lines against Orco (BL 31278) or mcherry (BL 35785) were used as controls, and UAP56 RNAis were VDRC 22557 (#1), BL 33666 (#2). Other dsRNA lines were Mical (VDRC 20672) and tsr (VDRC 110599). All dsRNAs were coexpressed with UAS-dcr2 (Dietzl et al., 2007) except in the MICAL-MS2 experiment in Figure 4. For MARCM, the P-element insertion *uap56^k11511^* (BL11043) was recombined on FRT40A. Other fly lines were *ppk-EB1::GFP* (Arthur et al., 2015) and act-Cas9, lig4^-/-^ (Zhang et al., 2014).

### Cloning and transgenes

C-terminally GFP-tagged UAP56 was cloned into pUAST attB, and ATPase-deficient UAP56::GFP E194Q was generated by PCR mutagenesis. The corresponding plasmids were injected into flies harboring the 86Fb insertion site. The non-cleavable ubiquitin reporter Ub-VV-GFP was cloned into pUAST attB and introduced into flies harboring VK37 or VK16 insertion sites, and a recombinant with both insertions was used. Ref1 was cloned into pENTR and subsequently into the destination vector pUAST myc rfa using a TOPO cloning strategy. In order to generate *MICAL-MS2_6_* flies, we generated a specific sgRNA (AGTAGAGAGCCTAACTAATA) targeting the the Mical 3’UTR in the pCFD3 vector. We then cloned tandem copies of the MS2 sequence (using pSL-MS2-6x (Addgene #27118) as a template) using the primers GGACGGATCCGCTTCTCCCATATTAGTTAGG (forward) and GGACCTCGAGAATGAACCCGGGAATACTG (reverse) and flanked them with 1.5 kb homology arms corresponding to sequences in the Mical CDS and 3’UTR. Primers for the flanking regions were: GGACGAATTCGTTTATGGCTTAAGTACTTGGC and GGACGGATCCGCTTCTCCCATATTAGTTAGG (left arm), and GGACCTCGAGGCGCAAGGTTAATGTAAAAACAG and GGACGCGGCCGCAAACACATCTCGTGACAACC (right arm). This homology donor was cloned into pBS SK(-), and the PAM motif for the sgRNA was mutated using PCR. The sgRNA and homology donor plasmids were then co-injected into *actin-Cas9, lig4^-/-^* embryos. Male injectants were crossed back, and male offspring was screened for the presence of MS2 by PCR. Positive lines were verified by PCR and sequencing, and two viable lines were chosen for experiments, both of which carried six copies of the MS2 sequences as assessed by sequencing.

### Dissection, Microscopy and Live imaging

For analyses of dendrite pruning phenotypes at 18 h APF, animals were dissected out of the pupal case and dorsal ddaC c4da neurons in segments A2 - A5 were imaged live on a Zeiss LSM710 confocal microscope with a 20x Plan Apochromat water objective (1.0 NA). EB1::GFP imaging was performed on a Zeiss LSM880 microscope using a 40x Plan Apochromat FCS M27 (1.2 NA) oil objective. Consecutive images of a single plane were taken every second for 1 - 2 minutes. For analyses of presynapse pruning, pupae were dissected out of the pupal case at 24 h APF and dissected ventrally. The brain and ventral nerve cord were removed and analyzed live using a Zeiss LSM710 confocal microscope with a 40x C-Apochromat water objective (1.1 NA). UAP56::GFP localization and the Ub-VV-GFP reporter fluorescence were assessed with a 40x C-Apochromat water objective (1.1 NA) at 3x zoom. For Mical mRNA analysis, UAS-MCP::RFP was expressed under *ppk-GAL4* in a *MICAL-MS2_6_* background. Images were taken live in undissected animals at the indicated developmental timepoints on a LSM710 confocal microscope using either a 40x C-Apochromat water objective (1.1 NA) or a 63x Plan-Apochromat oil objective (1.4 NA), and MCP::RFP dots were counted manually. All processing was done in Fiji (Schindelin *et al*, 2012) using the plug-in Image Stabilizer for EB1::GFP comet analysis.

### Immunofluorescence

Antibodies against Mical (Rode et al., 2018) (1:500) and Sox14 (Ritter and Beckstead, 2010) (1:30) were used as described (Rumpf et al., 2014, Herzmann et al., 2017). Appropriately staged pupae were dissected, and body wall filets were fixed in 4% formaldehyde for 20 minutes, washed in PBS with 0.3 % Triton X-100 and blocked in PBS with 0.3 % Triton X-100 and 10% goat serum. Antibody incubations were over night.

### Immunoprecipitation and Western blot

pUAST expression plasmids encoding wild type or mutant UAP56::GFP and myc-Ref1 were cotransfected into S2 cells with Actin5C-GAL4. After 72 hours, cells were harvested in ice-cold PBS and lysed for 30 minutes in lysis buffer (100 mM NaCl, 5 mM Mg(OAc)_2_, 5 % glycerol, 50 mM Tris/HCl pH 7.4, 1 % Triton X-100, 1x complete protease inhibitors). Cleared lysates were precipitated with rabbit anti-GFP antibodies (Invitrogen A11222) bound to protein A sepharose for two hours. After three washes in lysis buffer, precipitates were dissolved in SDS sample buffer at 95 °C for 5 minutes, run on an 8 % polyacrylamide gel and blotted with antibodies against myc (9E10) or GFP (JLA-8).

### Quantification and statistical analyses

Phenotypic penetrance was assessed by counting the number of neurons with dendrites still attached to the soma. Here, significance was determined using a categorical two-tailed Fisher’s exact test (graphpad.com). The number of primary and secondary dendrites attached to the soma (a measure of phenotypic strength) was counted manually and compared using the Wilcoxon Mann Whitney test (Marx et al., 2016). Ub-VV-GFP fluorescence intensity was taken after background subtraction and compared using student’s t-test. EB1 comet directionality across samples was compared using Fisher’s exact test, and comet speed was compared with Wilcoxon’s test using Prism software. The number of cytoplasmic MCP dots was counted manually, and compared using a t-test. To assess expression levels of Mical protein, Mical staining intensity in c4da neurons was normalized to its expression in unaffected neighboring c1da neurons. Data were then compared using Wilcoxon’s test. The number of synaptic Brp puncta per segment was determined from binarized confocal image stacks using the Fiji plugin SynQuant (Wang et al., 2020). Significance was determined in Prism using student’s t-test or ANOVA depending on normal distribution.

## Acknowledgments

We thank C. Klämbt for support, S. Sigrist and J. Wildonger for fly lines, the Bloomington and VDRC stock centers and Addgene for fly lines and reagents. SMF, SRo and MD are members of the SFB1348 and CiM graduate schools, respectively. This work was supported by the DFG Excellence Cluster “Cells in Motion” (EXC1003) and DFG grants RU1673/6-1 and RU/1673/10-1 to SR. The authors declare no competing financial interests.

## Author Contributions

SMF performed synapse experiments, UG and SRu performed most dendrite and imaging experiments. SRo performed the inital RNAi screen, MD performed the microtubule analysis. UG and SRu generated reagents. All authors contributed to experimental design. SRu analyzed data and wrote the manuscript with input from all other authors.

## Supplementary Figures

**Figure S1.**
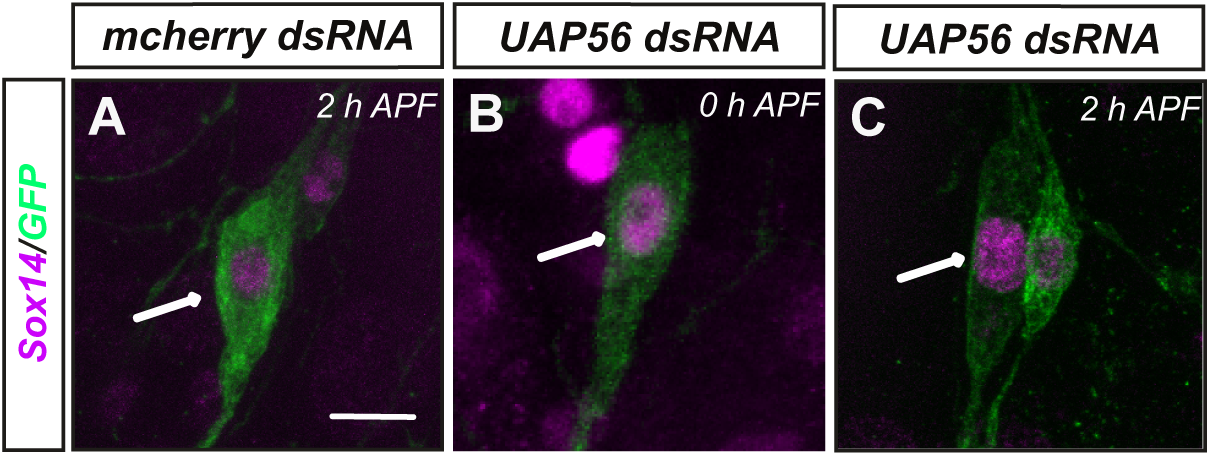
Sox14 expression in c4da neurons lacking UAP56. C4da neurons were labeled by *CD8::GFP* under the control of *ppk-GAL4* and Sox14 was detected by immunofluorescence. **A** Control c4da neuron expressing Orco dsRNA at 2 h APF. **B** C4da neuron expressing UAP56 dsRNA at 0 h APF. **C** C4da neuron expressing UAP56 dsRNA at 2 h APF. The scale bars in **A** is 10 μm.

**Figure S2.**
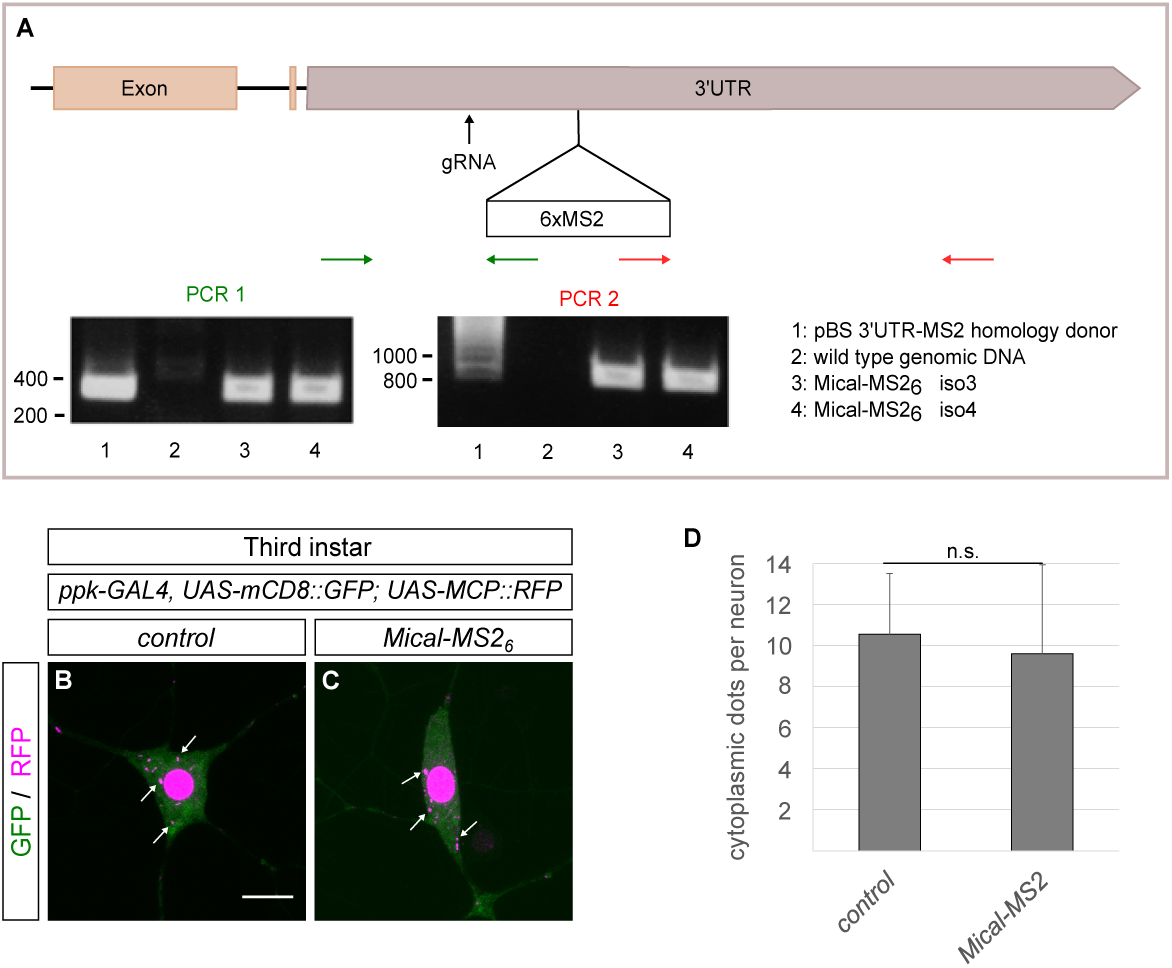
Generation and characterization of *Mical-MS2_6_*. **A** Strategy for generation of Mical-MS2 flies. The schematic shows the position of the sgRNA relative to the MS2 sequence insertion site and those of the PCR primers used to confirm successful insertion. The agarose gels show PCR confirmation of two positive fly lines with the primers indicated above. Running positions of size markers are indicated on the left of each gel. **B**, **C** Images of third instar larval c4da neuron somata expressing MCP::RFP under the control of *ppk-GAL4*. **B** C4da neuron expressing MCP::RFP in wild type background. **C** C4da neuron expressing MCP::RFP in Mical-MS2 background. **D** Quantification of the number of cytoplasmic MCP::RFP puncta in **B**, **C**. N=10 each, n. s., not significant, Student’s t-test. The scale bar in **B** is 10 μm.

## References

Arthur, A.L., Yang, S.Z., Abellaneda, A.M., Wildonger, J., 2015. Dendrite arborization requires the dynein cofactor NudE. J. Cell Sci. 128, 2191–2201.

Bertrand, E., Chartrand, P., Schaefer, M., Shenoy, S.M., Singer, R.H., Long, R.M., 1998. Localization of ASH1 mRNA particles in living yeast. Mol. Cell 2, 437–445.

Booth, K.T.A., Jangam, S.V., Chui, M.M.C., Treat, K., Graziani, L., Soldano, A., Ruan, Y., Wan-Hei Hui J., White, K., Christensen, C.K., Lynnes, T., Yamamoto, S., Kanca, O., Tsang, M.H.Y., Lynch, S.A., Mullegama, S.V., Batista, J., Iancu, D., Joss, S.K., Wong, S.Y.Y., Mak, C.C.Y., Kwong, A.K.Y., Bellen, H.J., Conboy, E., Sanges, R., Leung, A.Y., Wangler, M.F., Chung, B.H.Y., Vetrini, F., 2025. De novo and inherited variants in DDX39B cause a novel neurodevelopmental syndrome. Brain awaf035.

Davies, M., Sanal, N., Wolterhoff, N., Gigengack, U., Shen, Y., Hahn, I., Rumpf, S. 2025. Spectraplakin cooperates with noncentrosomal microtubule regulators to orient dendritic microtubules in Drosophila. biorxiv 2025.06.27.661943.

Dietzl, G., Chen, D., Schnorrer, F., Su, K.-C., Barinova, Y., Fellner, M., Gasser, B., Kinsey, K., Oppel, S., Scheiblauer, S., Couto, A., Marra, V., Keleman, K., Dickson, B.J., 2007. A genome-wide transgenic RNAi library for conditional gene inactivation in Drosophila. Nature 448, 151–156.

Grueber, W.B., Ye, B., Yang, C.-H., Younger, S., Borden, K., Jan, L.Y., Jan, Y.-N., 2007. Projections of Drosophila multidendritic neurons in the central nervous system: links with peripheral dendrite morphology. Development 134, 55–64.

Han, C., Song, Y., Xiao, H., Wang, D., Franc, N.C., Jan, L.Y., Jan, Y.-N., 2014. Epidermal cells are the primary phagocytes in the fragmentation and clearance of degenerating dendrites in Drosophila. Neuron 81, 544–560.

Herzmann, S., Gotzelmann, I., Reekers, L.-F., Rumpf, S., 2018. Spatial regulation of microtubule disruption during dendrite pruning in Drosophila. Development 145, pii: dev156950.

Herzmann, S., Krumkamp, R., Rode, S., Kintrup, C., Rumpf, S., 2017. PAR-1 promotes microtubule breakdown during dendrite pruning in Drosophila. EMBO J 36, 1981–1991.

Kirilly, D., Gu, Y., Huang, Y., Wu, Z., Bashirullah, A., Low, B.C., Kolodkin, A.L., Wang, H., Yu, F., 2009. A genetic pathway composed of Sox14 and Mical governs severing of dendrites during pruning. Nat. Neurosci. 12, 1497–1505.

Krämer, R., Wolterhoff, N., Galic, M., Rumpf, S., 2023. Developmental pruning of sensory neurites by mechanical tearing in Drosophila. J. Cell Biol. 222.

Kuo, C.T., Jan, L.Y., Jan, Y.N., 2005. Dendrite-specific remodeling of Drosophila sensory neurons requires matrix metalloproteases, ubiquitin-proteasome, and ecdysone signaling. Proc. Natl. Acad. Sci. U.S.A. 102, 15230–15235.

Meader, C.I., Kim JI, Liang X, Kaganovsky K, Shen A, Li Q, Li Z, Wang S, Xu XZS, Li JB, Xiang YK, Ding JB, Shen K., 2018. The THO Complex Coordinates Transcripts for Synapse Development and Dopamine Neuron Survival. Cell 174:1436–1449.e20.

Marzano, M., Herzmann, S., Elsbroek, L., Sanal, N., Tarbashevich, K., Raz, E., Krahn, M.P., Rumpf, S., 2021. AMPK adapts metabolism to developmental energy requirement during dendrite pruning in Drosophila. Cell Rep. 37, 110024.

Matsubara, D., Horiuchi, S.-Y., Shimono, K., Usui, T., Uemura, T., 2011. The seven-pass transmembrane cadherin Flamingo controls dendritic self-avoidance via its binding to a LIM domain protein, Espinas, in Drosophila sensory neurons. Genes Dev. 25, 1982–1996.

Park, J.W., Parisky, K., Celotto, A.M., Reenan, R.A., Graveley, B.R., 2004. Identification of alternative splicing regulators by RNA interference in Drosophila. Proc. Natl. Acad. Sci. U.S.A. 101, 15974–15979.

Rajan, S., Yoon, J., Wu, H., Srapyan, S., Baskar, R., Ahmed, G., Yang, T., Grintsevich, E.E., Reisler, E., Terman, J.R., 2023. Disassembly of bundled F-actin and cellular remodeling via an interplay of Mical, cofilin, and F-actin crosslinkers. Proc. Natl. Acad. Sci. U.S.A. 120, e2309955120.

Riccomagno, M.M., Kolodkin, A.L., 2015. Sculpting neural circuits by axon and dendrite pruning. Annu. Rev. Cell Dev. Biol. 31, 779–805.

Ritter, A.R., Beckstead, R.B., 2010. Sox14 is required for transcriptional and developmental responses to 20-hydroxyecdysone at the onset of drosophila metamorphosis. Dev. Dyn. 239, 2685–2694.

Rode, S., Ohm, H., Anhauser, L., Wagner, M., Rosing, M., Deng, X., Sin, O., Leidel, S.A., Storkebaum, E., Rentmeister, A., Zhu, S., Rumpf, S., 2018. Differential Requirement for Translation Initiation Factor Pathways during Ecdysone-Dependent Neuronal Remodeling in Drosophila. Cell Rep. 24, 2287–2299.e4.

Rode, S., Ohm, H., Zipfel, J., Rumpf, S., 2017. The spliceosome-associated protein Mfap1 binds to VCP in Drosophila. PLoS One 12, e0183733.

Rumpf, S., Bagley, J.A., Thompson-Peer, K.L., Zhu, S., Gorczyca, D., Beckstead, R.B., Jan, L.Y., Jan, Y.N., 2014. Drosophila Valosin-Containing Protein is required for dendrite pruning through a regulatory role in mRNA metabolism. Proc. Natl. Acad. Sci. U.S.A. 111, 7331–7336.

Rumpf, S., Lee, S.B., Jan, L.Y., Jan, Y.N., 2011. Neuronal remodeling and apoptosis require VCP-dependent degradation of the apoptosis inhibitor DIAP1. Development 138, 1153– 1160.

Rumpf, S., Rode, S., Herzmann, S., Krumkamp, R. 2017. Mechanisms of Neurite Pruning. Neuroforum 23, 19–25.

Rumpf, S., Wolterhoff, N., Herzmann, S., 2019. Functions of Microtubule Disassembly during Neurite Pruning. Trends Cell Biol. 29, 291–297.

Rust, M. B., 2015. ADF/cofilin: a crucial regulator of synapse physiology and behavior. Cell. Mol. Life Sci. 72, 3521–3529.

Schindelin, J., Arganda-Carreras, I., Frise, E., Kaynig, V., Longair, M., Pietzsch, T., Preibisch, S., Rueden, C., Saalfeld, S., Schmid, B., Tinevez, J.-Y., White, D.J., Hartenstein, V., Eliceiri, K., Tomancak, P., Cardona, A., 2012. Fiji: an open-source platform for biological-image analysis. Nat. Methods 9, 676–682.

Terman, J.R., Mao, T., Pasterkamp, R.J., Yu, H.-H., Kolodkin, A.L., 2002. MICALs, a family of conserved flavoprotein oxidoreductases, function in plexin-mediated axonal repulsion. Cell 109, 887–900.

Wang, Y., Wang, C., Ranefall, P., Broussard, G. J., Wang, Y., Shi, G., Lyu, B., Wu, C.-T., Wang, Y., Tian, L., Yu, G., 2020. SynQuant: an automatic tool to quantify synapses from microscopy images. Bioinformatics 36, 1599–1606.

Williams, D.W., Truman, J.W., 2005. Cellular mechanisms of dendrite pruning in Drosophila: insights from in vivo time-lapse of remodeling dendritic arborizing sensory neurons. Development 132, 3631–3642.

Wolterhoff, N., Gigengack, U., Rumpf, S., 2020. PP2A phosphatase is required for dendrite pruning via actin regulation in Drosophila. EMBO Rep. 21, e48870.

Xie Y, Ren Y., 2019. Mechanisms of nuclear mRNA export: A structural perspective. Traffic 2019 20, 829–840

Zhang, X., Koolhaas, W.H., Schnorrer, F., 2014. A versatile two-step CRISPR- and RMCE-based strategy for efficient genome engineering in Drosophila. G3 4, 2409–2418.

